# A home-cage, video monitoring-based mouse frailty index detects age-associated morbidity in the absence of handler-induced stress

**DOI:** 10.1101/2022.07.19.500666

**Authors:** J. Graham Ruby, Paulo Ylagan, Andrea Di Francesco, José Zavala-Solorio, Robert Keyser, Owen Williams, Sarah Spock, Wenzhou Li, Nalien Vongtharangsy, Sandip Chatterjee, Cricket A. Sloan, Charles Ledogar, Veronica Kuiper, Janessa Kite, Marcelo Cosino, Paulyn Cha, Eleanor M. Karlsson

## Abstract

Frailty indexes provide quantitative measurements of non-specific health decline and are particularly useful as longitudinal monitors of pre-mortal morbidity in aging studies. For mouse studies, frailty assessments can be taken non-invasively, but they require handling and direct observation that is labor-intensive to the scientist and stress-inducing to the animal. Here, we implement, evaluate, and provide a digital frailty index composed entirely of computational analyses of home-cage video and compare it to manually obtained frailty scores in genetically diverse mice. We show that the frailty scores assigned by our digital index correlate with both manually obtained frailty scores and chronological age. Thus, we provide a tool for frailty assessment that reduces stress to the animal and can be collected consistently, at scale, without substantial labor cost.

## Introduction

For many organisms on the planet Earth, increasing age for an individual is accompanied by physiological deterioration and an increase in mortality hazard. At a population level, physiological decline with age can be measured using mortality statistics, and for many species, the risk of death increases exponentially with age (Jones et al, 2014). While useful for performing comparisons between species or populations, mortality provides only one datum per individual and therefore requires large cohorts and precludes longitudinal analysis of individuals.

In humans, the physiological decline that accompanies age can also be measured in terms of a “frailty index” (FI): a tally of health-related deficits that are clinically observable (Clegg et al, 2013). Many varieties of human FI exist, with variable focus on performance-based phenotypic assessment (e.g. Fried et al, 2001) or the accumulation of overt pathologies (e.g. Song et al, 2010) but consistently predict the likelihood of future disability and mortality (e.g. Rockwood et al, 1999). Due to their implicitly multifaceted nature, frailty assessments often require substantial time, cost, and expertise to administer. Many variants of human FI implementation attempt to reduce those practical constraints: among them, a version requiring minimal and non-expert examination (Rolfson et al, 2006); a version derived exclusively from blood-test measurements (Howlett et al, 2014); and a version based solely on retrospective analysis of electronic health records (Clegg et al, 2016).

The FI concept is also applied to mice in preclinical research (Parks et al, 2011). As with humans, alternative versions of a mouse FI have been implemented that reflect diverging priorities: some emphasize thoroughness and include laboratory tests (e.g. Kane et al, 2019); others attempt to maximize similarity to human FI (Liu et al, 2014). Most of these implementations involve subjective scoring by a technician, which poses two challenges to the practical application of FI. Firstly, the multitude of traits to be examined places substantial demand on the time of the researcher performing FI assessments. Secondly, substantial inter-rater variance has been documented. The sources of that variance remain debatable, but it can be exacerbated by variance across raters in years of education or direct experience (Feridooni et al, 2015; Kane et al, 2015; Howlett & Rockwood, 2015).

Consistency of mouse FI across raters has been mitigated through automation of analysis. For grimace – a commonly-included component of manual mouse FI’s that indicates the mouse’s experience of discomfort (Langford et al, 2010) – machine-learning (ML) assessments perform similarly to human raters when applied to images captured in tabletop cubicles (Tuttle et al, 2018). Parameters derived from the open-field assay (Walsh & Cummins, 1976), for which mouse behavior is often analyzed using computer vision (Seibenhener & Wooten, 2015), are often included in traditional frailty assessments (e.g. Parks et al, 2011). The convenience of the open-field paradigm for analysis of multiple traits has motivated the development of mouse FI versions that rely mostly (Whitehead et al, 2013) or entirely (Hession et al, 2021) on analytics of data from that assay. In addition to improved consistency/reliability, these cases greatly ease the practical application of FI by consolidating data collection to a single, low-labor assay.

The approaches to mouse frailty assessment described above vary in terms of their invasiveness, but they invariantly require handling and/or exposure to a non-standard environment in order to be performed. Handling and exposure to novel environments both cause stress in laboratory mice (Balcombe et al, 2004; Baran et al, 2022). The frailty indexes described above invariantly include parameters known to be modified by handling-related stress: posture, response to stimuli, exploratory behavior, and blood chemistry among them (Hurst & West, 2010; Ghosal et al, 2015; Gouveia & Hurst, 2017). Handling could also further stress already compromised or frail animals and push them towards humane endpoints, exacerbating survivor bias in aging studies. The recent development of systems capable of continuously monitoring laboratory mice in their home-cage environment (Lim et al, 2017; Baran et al, 2021) provides an avenue to refine longitudinal aging studies and avoid these stresses as confounders of frailty assessment and longevity. But that advancement requires the development of analytics capable of measuring frailty from home-cage video data.

Here, we implemented a digital frailty index (DFI) for mice based on the computational analysis of continuously-collected home-cage video footage. To evaluate its effectiveness, we performed a study involving over 200 mice in which video was collected longitudinally, in parallel with manual frailty indexes (MFI) collected at a massive scale, all by a single researcher. Our implementation of DFI correlated with both chronological age and MFI, the correlation between DFI and MFI was maintained when the age-related components of both measurements were regressed out. While the inclusion of additional parameters may enhance DFI value in the future, here we prove the feasibility, scalability and relevance of frailty assessment in mice through passive observation in a home-cage environment.

## Methods

### Mice, animal husbandry, and in vivo study design

For the main Frailty study, 228 Diversity outbred (J:DO) mice (138 female, 90 male; The Jackson Laboratory, Strain #009376) across eight aged cohorts were obtained from The Jackson Laboratory. Mice were aged to 6, 9, 12, 15, 19, 21, 25, 27 and 30 months at the beginning of the study period and weighed between 18g and 67g. Video collection and manual frailty assessments were each performed at three time points, at six week intervals. Females were group housed when not in video cages, and males were singly housed throughout the study. Mice were shipped from The Jackson Laboratory (Bar Harbor, ME) and acclimated for at least 2 weeks. Census data on these mice are provided in Supplemental Table 1.

All mice were housed in solid-bottom 100% PET plastic, BPA-Free IVC cages (Innovive). All cages and bedding were irradiated prior to use. Mice were housed on 1/8-in. corn cob bedding (Innovive, San Diego, CA), received acidified (pH 2.5 to 3.0) reverse osmosis–purified water from water bottles (Innovive, San Diego, CA), and were fed irradiated diet chow (LabDiet Pico 5L0D, Purina, St. Louis, MO). Mice received 8g shredded paper nesting material and a paper hut enrichment (Enviro-Dri and Shephard Shack, Shephard Specialty Papers, Watertown, TN) while in their non-video cages. Mice were provided two cotton nesting squares (NES 3600, Ancare, Bellmore, NY) for enrichment, along with a climbing ladder and running wheel (Vium) while in video cages. All mice were housed under a 12:12-h light:dark cycle at a density of 1 to 4 mice per cage in a temperature-controlled vivarium, in compliance with the Guide for the Care and Use of Laboratory Animals. Male mice were singly housed throughout the study, while female mice were single housed in video cages (1 week out of every 6 weeks for a total of 18 weeks). Animal cages were changed every 2 weeks within a cage change station (NuAire, Plymouth, MN). Mice were transferred between cages using red transparent acrylic tunnels (Bio-Serv, NJ) or by cupping technique.

Manual frailty was assessed longitudinally by the same experimenter as described in the original 31-item mouse clinical FI (Whitehead 2014), omitting the two parameters that rely on population-specific statistics (body weight and temperature). Mice were allowed to acclimatize to the testing room for 30-45 minutes before testing. The observation was carried out on an open bench at the same time of the day, between 9-11am. Mice were scored 0, 0.5, or 1 based on the severity of deficit they showed in each of the 29 items, with 0 representing no sign of deficit, 0.5 mild deficit and 1 severe deficit. Those 29 items were: alopecia, loss of fur color, dermatitis, loss of whiskers, coat condition, breathing rate/depth, mouse grimace scale, piloerection, tumors, distended abdomen, kyphosis, gait disorders, tremor, vestibular disturbance, tail stiffening, cataracts, corneal opacity, eye discharge/swelling, microphthalmia, nasal discharge, malocclusions, rectal prolapse, vaginal/uterine/penile prolapse, diarrhea, body condition score, forelimb grip strength, menace reflex, vision loss, and hearing loss. The overall manual frailty index (MFI) was taken as the average score across those 29 parameters. These data are provided in Supplemental Table 2.

Body weight was measured at the beginning of each test and body surface temperature was measured averaging three readings obtained with an infrared temperature probe directed at the abdomen. Because of the potential for genetic variance in the J:DO population to confound the interpretation of specific values for these parameters as frailty-related, they were not included in the MFI scores but are provided in Supplemental Table 2, along with other MFI scores.

Video collection was performed by placing cages with singly-housed mice into video camera-equipped racks for approximately one week. Video footage of mouse cages was streamed 24 hours a day, 7 days a week to a cloud-based data infrastructure. Video acquisition hardware and compatible cage furniture were purchased from Vium (https://www.vium.com). Data management infrastructure was created by the authors of this study.

Video data for model training and parameterization were collected separately, under similar conditions as for the main Frailty study (described above). Those data included Diversity outbred (J:DO) mice, aged to 19 months (n = 40), 26 months (n = 37), and 32 months (n = 31) and monitored for one week; and C57BL/6J mice, aged to 7, 19, and 35 months (n = 10 per age cohort) and monitored for three weeks. All videos were collected at 864 pixel (width) by 648 pixel (height) resolution, approximately 24 frames per second, from cameras mounted in consistently fixed positions versus the cage and furniture (e.g. Fig. 1A).

**Figure 1:**
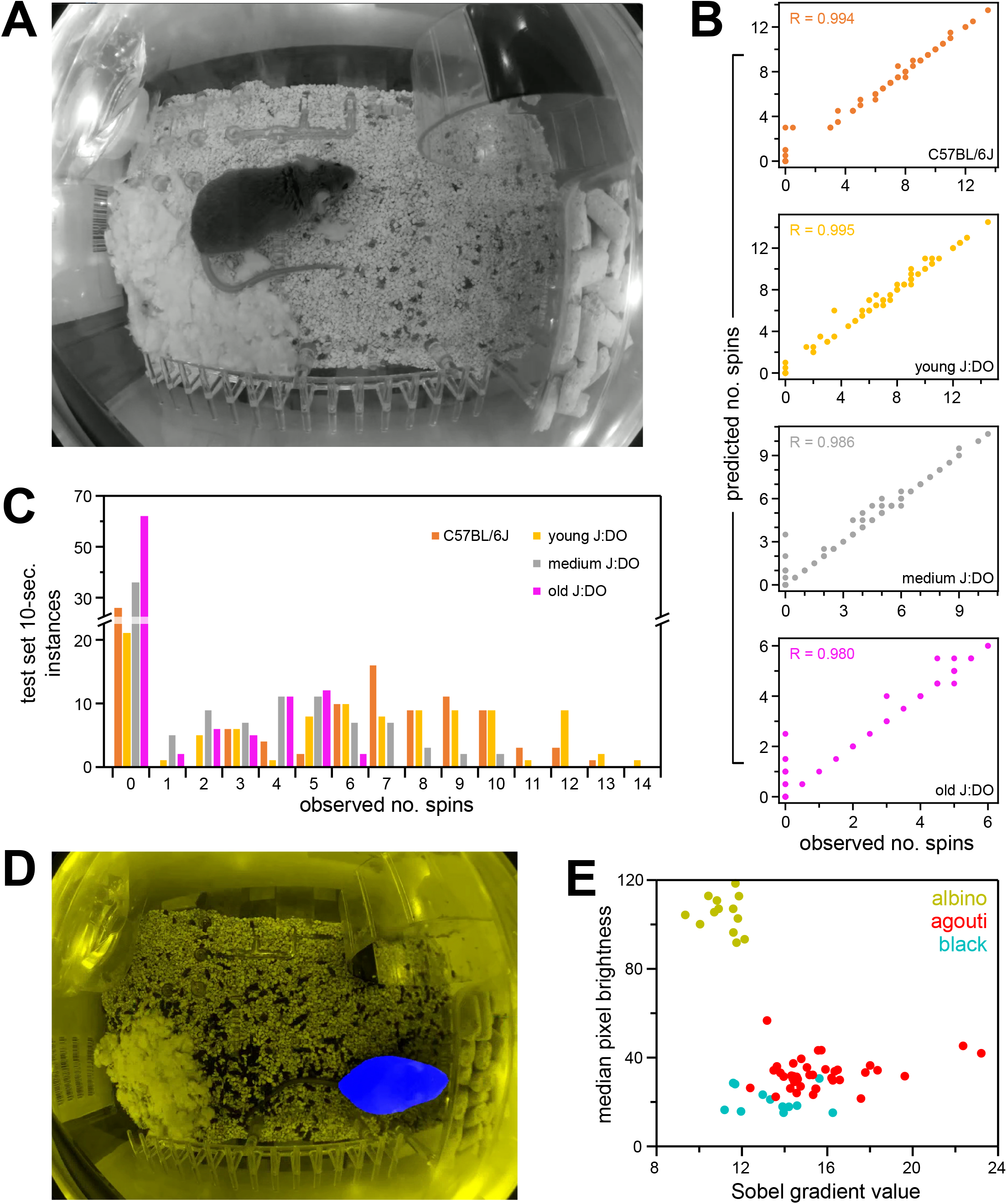
Example video stills and methodological details. A) An example frame of video footage, typical of nighttime video collected in this study. Nighttime illumination was near-IR, resulting in grayscale images. Daytime images were similar, but in RGB color. B) Performance of the wheel-spin detection pipeline versus test data. For each of four different cohorts of mice, 100 10-second video clips were manually annotated for the number of wheel spins that occurred, in increments of half-spins, based on appearance and disappearance of the black marker. Those values are correlated against the number of spins annotated by the ML pipeline across the same time segment. The four cohorts included footage of either C57BL/6J mice or young, medium-age, or old J:DO mice. See Methods for details. C) The distributions of wheel-spin counts across 10-second clips from the four manually annotated test sets from panel (B). D) An example of a modified input image for body-weight prediction error-detection models, with the blue channel modified to indicate masked (blue) versus unmasked (yellow) output from the mouse semantic segmentation model. See Methods for details. E) Sobel values versus median pixel brightness for training-set J:DO mice, colored according to human assignment of coat color.

All research was performed as part of Calico Life Sciences LLC AAALAC-accredited animal care and use program. All research and animal use in this study was approved by the Calico Institutional Animal Care and Use Committee (IACUC).

### Calculation of Digital Frailty Index (DFI) score

Eight frailty parameters were scored: average per-day distance run on the wheel; gait speed on the wheel; gait speed on the floor of the cage; circadian distribution of wheel-running activity; circadian distribution of movement on the floor of the cage; rate of change in body weight; coat quality; and average displacement of nesting material. The derivation of each measure from raw video footage is described below, followed by the parameterization rules for converting each measure to a frailty score component.

### Wheel-derived DFI parameters

Three parameters derived from analysis of wheel spins: average per-day distance run on the wheel; gait speed on the wheel; and circadian distribution of wheel-running activity. Wheel-running activity was monitored using an ML pipeline that identified the timing of individual rotations of the running wheel using a black stripe on the wheel’s surface (Fig. 1A). In our video cages, the running wheel always appeared in the upper-right quadrant of the image, so all ML models in this pipeline were trained on and applied to only that quadrant of each video frame.

The pipeline first applied two independent, two-channel image-segmentation models: the first to draw a mask covering the open face of the running wheel, the second to draw a mask covering the black stripe. Those masks were used to construct a synthetic image, with the wheel-face mask displayed in the blue channel and the black-stripe mask displayed in the green channel. Overlap of the two masks, which occurred when either the stripe was under but still visible through the translucent-plastic wheel or when a black mouse was confused as a stripe, therefore appeared as cyan in the synthetic image. That synthetic image was provided as input to an image-classification ML model, which returned probabilities for each of two classes, corresponding to the black stripe being on the “top” or “bottom” of the wheel. Those probabilities were used as emission probabilities for those two respective states, and a two-state Hidden Markov Model (HMM) was parsed across those emissions using the Viterbi algorithm (Viterbi, 1967) across all frames of each full 10-minute video. From the resulting parse, every full transition cycle (from “top” to “bottom” and back to “top”, or vice-versa, depending on the initial state for that video) was recorded as one spin of the wheel, across a length of time determined by the video frame rate and the number of frames traversed through the spin.

The two image-segmentation models described above, for the wheel’s face and marker, were each trained using the tool provided by the “image-segmentation-keras” code base (https://github.com/divamgupta/image-segmentation-keras commit f04852d from September 6, 2019), specifying the “vgg_unet” model architecture (Simonyan, 2014), with input widths and heights of 128 and 96, respectively. These models were trained on and applied to only the upper-right quadrant of the images, which consistently contained the entire wheel. The classification model described above was a re-trained version of the TF-Slim implementation of mobilenet_v1 with a depth multiplier of 1.0 and an input size of 224 by 224 (Howard et al, 2017). The wheel-position HMM had initial state probabilities of 0.325 for “top” and 0.675 for “bottom”. Transition probabilities were set to ⅔ when returning to the same state and ⅓ when changing state.

The wheel-spin ML pipeline was trained on frames taken from video of C57BL/6J mice. It was tested on both an independent set of C57BL/6J mice and video from three cohorts of differently-aged J:DO mice, using manually spin counts for 100 10-second clips from each cohort. Correlation coefficients (Pearson’s R) between the model- and manual-derived spin counts were 0.994 for the C57BL/6J cohort and 0.995 (young), 0.986 (middle), and 0.980 (old) across the three J:DO cohorts (Fig. 1B). Across all four test cohorts, accuracy increased with the mean number of manually-counted spins per clip – 5.73 spins/clip for the C57BL/6J cohort and 1.52 spins/clip (old), 3.15 spins/clip (middle), and 6.08 spins/clip (young) for the J:DO cohorts (Fig. 1C). By inspection: most errors were single frames at the beginning or end of the clip for which the status of transition to a new state (“top” or “bottom”) was ambiguous.

To calculate the average per-day distance run on the wheel: for each ten-minute video, frame-by-frame parsed HMM states were analyzed to count wheel spins by counting the number of instances of the state changing (between the marker being on the “top” or “bottom” of the wheel) and dividing by two. The wheel-spin sums from all videos within each designated DFI measurement period (approx. one week) were summed and divided by the total length of footage from that DFI measurement period, in days.

To calculate gait speed on the wheel: for each ten-minute video, frame-by-frame parsed HMM states were analyzed to identify the number of frames that elapsed for each complete spin (number of frames divided by frames-per-second for the video). For all complete spins observed across the designated DFI measurement period, the median length of time *t*_*sp_med*_ was used to define the typical gait speed (1 / *t*_*sp_med*_).

To calculate the circadian distribution of wheel-running activity: distance in wheel spins was determined for each 10-minute video, as was done for total distance. For each designated DFI measurement period, day versus night periods were defined as alternating non-overlapping, 12-hour blocks of clock time, and their phase was allowed to vary by 10-minute increments across the full 24-hour cycle. For each phase, the average activities across “day” and “night” videos were calculated, and the differences between the two were recorded. The maximum absolute value from those differences was used to define circadian activity, normalized to twice the weighted average across all “day” and “night” values.

### Floor-of-cage movement DFI parameters

Two DFI parameters derived from analysis of movement across the floor of the cage: gait speed on the floor of the cage; and circadian distribution of movement on the floor of the cage. Tracking of the mouse as it moved about the cage provided the foundational data for multiple parameters. This tracking was achieved using an object-detection model, trained by transfer learning using the COCO-trained ssd_mobilnet_v1 model (downloaded from http://download.tensorflow.org/models/object_detection/ssd_mobilenet_v1_coco_11_06_2017.tar.gz) (Liu et al, 2016; Howard et al, 2017) using images taken from video feeds. Since all video footage was taken of singly-housed mice for this study, only the most confidently predicted box was used from each frame, and it was used regardless of its confidence score.

To calculate gait speed on the cage floor, the position of the mouse was determined for each frame of video using the center point of the highest-confidence box drawn by the mouse-detector model. For each frame in a 10-minute video, a refined position estimate was calculated using the mean position of the mouse across 15 adjacent frames (this truncated seven frames each from the beginning and end of each video). Using those refined positions, movement between each pair of frames was calculated as the linear distance between the positions in the two frames, in units of pixel length, using the Pythagorean theorem. Per-second distances were calculated for each non-overlapping second of video, using the integer of the video-encoded frame rate (fps), multiplied by the ratio of the float-encoded fps to its integer value (to compensate for under-estimates of distance due to frame-unit truncation). Gait speed was calculated as the weighted average of distance-per-second values, weighted by the distance-per-second value, which was mathematically equivalent to calculating the average gait speed per unit of distance covered. These weighted averages were taken across all per-second estimates available across a day. Overall gait-speed estimates for a mouse across a frailty-measuring time segment were calculated as the weighted averages of all per-day estimates from the time interval, weighted to the total distance traveled per day.

Circadian movement on the floor of the cage was calculated using the same distance-traveled values described above for gait speed. The same method was applied as for the circadian distribution of wheel-running activity, substituting distance-traveled for wheel-spins.

### Body weight change DFI parameter

For the estimation of body weight and assessment of coat condition, pixels overlapping the mouse (i.e. mouse masks) were identified through a two-step process, performed on each frame of video. First, the most confidently predicted bounding-box from the mouse-detector model described above was used to isolate a subset of the frame image, corresponding to the area of the bounding box extended by 20% along each dimension (10% extension in each direction). Second, that sub-image, which constituted a zoom-in on the mouse, was fed as input to an image segmentation model that would provide a mask of the mouse as output. That segmentation model was trained using “image-segmentation-keras” as described above, again specifying the “vgg_unet” model architecture (Simonyan, 2014), with input width and height both set to 256. That model was trained & tested using similarly-boxed images of J:DO mice and achieved a mean IoU of 0.905 (SD 0.050) across 296 test images.

For the per-frame body weight estimate, the area of the mouse mask was normalized to remove fish-eye lens effects, based on each pixel’s position in the original image. The area of each pixel *a* was up-weighted using the following equations, where *x*_*eq*_ and *y*_*eq*_ are the x and y coordinates of the pixel in the 864 pixel (width) by 648 pixel (height) image, relative to the center of the image, and the focal length variable *f* was assigned a value of 415.0:

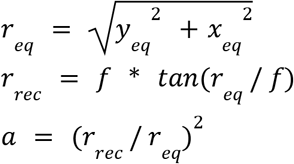

That measurement of area of the mouse in the image, in pixels, was then converted to voxels by accounting for the ratio of cross-sectional area to volume for a sphere (π * r^2^ versus 4/3 * π * r^3^), by raising the pixel area to the power of 1.5 (the constant 4/3 was ignored). Based on regression with training data (description of training data set to follow), the resulting volume in voxels was converted to body weight in grams using the conversion factor: 1.309 * 10^−5^ voxels per gram.

For each body weight estimate, an image classification model was applied to identify and compensate for the largest-error estimates. This model was trained empirically by applying the above model to the training data set and categorizing images based on the magnitude and direction of the estimates’ errors. Three classes were defined, separating training-set images based on errors >1.8-fold from the true value (too low for class “a”; within that range for class “b”; too high for class “c”). A classification model was trained using transfer learning, starting from EfficientNet-B0 (Tan & Le, 2019; downloaded from https://tfhub.dev/google/efficientnet/b0/classification/1). Input images for that model were modified versions of the full original image, with the green and red channels of each pixel set to its grayscale value (average brightness value across the three color channels) and the blue value set to maximum brightness (255) in masked pixels and minimum brightness (0) in un-masked pixels (e.g. Fig. 1D). That modified image was classified as likely to be either an over-estimate, under-estimate, or appropriate estimate of body weight. Body weights from over-estimate images were multiplied by 0.4, and body weights from under-estimate images were multiplied by 2.174, based on training data error averages in these categories.

The same modified image was used by a second classification model to assign a reliability weight to the body weight estimate. Two classes were defined: “mouse” or “other”. Training and input images were colored in the same manner as input images for the estimate-error classification model above, with the segmented “mouse” pixels differentially colored. Training images were manually sorted based on a human rater’s opinion of whether the mask substantially covered the mouse and not substantially other parts of the image. The classification model was again trained using transfer learning, starting from EfficientNet-B0 (Tan & Le, 2019). In the DFI-scoring pipeline, images classified as “other” were assigned reliability weights of zero. Images classified as “mouse” were assigned a reliability weight *w*_*rel*_ derived from the confidence score *c* assigned to that class by the model: *w*_*rel*_ = 2*(*c* - 0.5). Reliability weights were recorded along with their corresponding body weight estimates for use in weighted regression or mean estimates, performed later.

Body-weight prediction models were trained and evaluated using an independent data set of J:DO mouse videos that were collected within a day of a manually-ascertained body weight measurement. From a pool of 2,246 weight measurements with matched video, 14,321 frames corresponding to 974 ground-truth weights (37.6g mean; 8.6g SD; 19.18g minimum, 71.24 maximum weights) were selected for training, with frames corresponding to the same weight measurement spaced at least 1.5 hours apart. For testing, 230 weight measurements were selected that did not overlap with the training set, each of which had at least 50 ten-minute video files captured on the same date as the weight measurement. For evaluation, body weights were estimated from the first frame of each ten-minute video on the calendar day (up to 144 predictions), and the weighted mean of the predictions from each mouse & day was compared to the ground-truth measurement for that mouse on that day.

For DFI, the rate of change of body weight was calculated across each designated DFI measurement period. Video frames were sampled once per minute, and estimates and accompanying weights were ascertained as described above for each video frame image. Rates of body-weight change were estimated using weighted linear regression with the time at which the video frame was recorded as the independent variable.

### Coat condition and nest movement DFI parameters

For the assessment of coat condition, the mouse detector model used above in the first step of mouse identification was again used to define a zoomed-in image of the mouse (as above: extended by 20% along each dimension). Sobel gradients (Sobel & Feldman, 1968) at each pixel were calculated using the OpenCV “Sobel” function (Bradski, 2000), along the x-axis (*s*_*x*_) and y-axis (*s*_*y*_) separately, then calculating the pixel value *s*_*p*_ as √(*s*_*x*_ ^2^ + *s*_*y*_^2^). The overall Sobel value *s*_*m*_ was calculated as the mean of *s*_*p*_ values for all pixels overlapping the mouse mask, defined as described for body weight. The brightness of the mouse’s coat *c*_*br*_ was also calculated as the median pixel intensity for all pixels overlapping the mouse mask. For both *s*_*m*_ and *c*_*br*_, the overall value for an animal, for a designated DFI measurement period, was taken as the mean of all estimates collected from across that period, with one frame sampled per minute. Due to systematic bias of the intensity of Sobel gradients across mice of different coat colors (Fig. 1E), the final *s*_*m*_ value was adjusted based on coat color, as ascertained by coat brightness. For mice with an average *c*_*br*_ less than 30 (black mice), the *s*_*m*_ value was increased by 2; and for mice with an average *c*_*br*_ greater than 70 (albino mice), the *s*_*m*_ value was increased by 4. For the remaining mice (agouti), the *s*_*m*_ value was not adjusted.

For the assessment of nest movement, a semantic segmentation model was trained to identify nest material (cotton in the video cages). Nest material was manually labeled for training and test-set images from an independent video data set. The segmentation model was trained using “image-segmentation-keras” as described above, again specifying the “vgg_unet” model architecture (Simonyan, 2014), with input width set to 768 and input height set to 512. That model achieved mean IoU’s of approximately 0.75 across two separately-annotated test sets (both manually annotated, months apart but by the same rater). The position of nest material was taken for each frame as the mean position of masked pixels. Positions were estimated for each ten-minute video from across each designated DFI measurement period: for each video, the position was calculated as the average of positions from frames sampled once per minute across the video. Position was specified in terms of the pixel grid of the source images. Displacement was measured between time-adjacent videos, in pixels per ten-minute interval. In cases of a missing video file, displacement rate was calculated between nearest-time-adjacent videos, with the displacement rate still calculated per ten-minute interval. The output value was the average of all displacement-rate values obtained across the designated DFI measurement period.

### Parameterization of measurements into frailty values

For each of the eight components of DFI, the measured value was converted into a frailty score, meant to indicate observation of an exceptional value that reflects an abnormal, pathological state. Threshold values were determined *ad hoc*, with guidance provided by two age-stratified training sets (one of C57BL/6J mice, the other of J:DO mice; described above). To mimic the structure and concept of MFI and similar frailty indexes, each of the eight components was scored on a scale from zero to one, with zero indicating normal and one indicating frail behavior. In MFI, quantification of intermediate or ambiguous frailty was permitted, to a very limited extent, through the optional assignment of a score of 0.5. For DFI, threshold values were established for scores of either zero or one, and intermediate scores were linearly represented in terms of their relative proximity to those two thresholds. The overall DFI value was calculated as the average of the eight parameterized scores. Those component scores and the overall score are provided for each measurement of DFI in Supplemental Table 3.

For gait speed on the wheel, frailty values of zero and one were thresholded at 0.5 and 0.1 spins per second, respectively. For gait speed on the floor of the cage, frailty values of zero and one were thresholded at 60 and 20 pixel-lengths per second, respectively. For circadian behavior on both the wheel and the floor of the cage, frailty values of zero and one were thresholded at circadian ratios of 0.6 and 0.2, respectively. For total wheel-running distance, frailty values of zero and one were thresholded at 5,000 and zero spins per day, respectively. For nest movement, frailty values of zero and one were thresholded at 7.5 and 2.5 pixel-lengths per ten-minute interval, respectively. For coat quality, frailty values of zero and one were thresholded at Sobel values of 20 and 25, respectively. For body weight change, frailty values of zero and one were thresholded for the absolute value of the per-day weight change, at 1 and 2 grams per day, respectively. For each component of each DFI measurement, the pre-parameterized phenotype values are provided in Supplemental Table 4.

### Data properties and statistical analyses

For analysis of MFI versus age, all recorded and properly formatted MFI measurements were used, with age calculated as the difference between the animal’s cohorts’ date of birth and the date of the MFI assay.

For video monitoring, footage was streamed into cloud storage from a camera-enabled cage rack, with one camera per cage position. Data were organized by the device ID of the camera that collected it, and by date and time of acquisition. An electronic system maintained records of which mouse occupied which cage position at what times and those records were used to define DFI measurement period. For each DFI measurement, a mouse was recorded in a video cage for approximately one week, singly housed, one week after MFI measurement. For analysis of DFI, incomplete days of video at the beginning and end of the designated DFI measurement period were excluded from the analysis. Only DFI periods with at least two full days of observation were included; for observation periods with more than six full days of observation, only the first six full days were considered.

Each DFI measurement was assigned to the first full day of video recording as the date of measurement. For analysis of DFI versus age, all DFI measurements meeting the criteria above were used, with age calculated as the difference between the animal’s cohorts’ date of birth and the first date of the DFI measurement date. A table of the measurement intervals that is compliant with the input requirements for the software system is provided in Supplemental Table 5.

For the comparison of MFI versus DFI, measurement pairs were ascertained by seeking, for each DFI measurement, the MFI measurement from the same mouse with the closest date of acquisition. Only pairs of data collected fewer than eight days apart were considered, with the DFI assay date corresponding to the beginning of DFI video acquisition. A table of qualifying MFI/DFI pairs is provided in Supplemental Table 6. For comparisons between age-normalized MFI and age-normalized DFI, linear regression was performed between age and the subset MFI or DFI data for which paired data had been found. The slopes calculated from those two regressions were used to calculate residuals of each datum versus its respective regression against age, and the MFI residuals were regressed against and correlated with the DFI residuals.

All regressions and correlations were calculated using the “scipy.stats.linregress” function from Scipy (Virtanen et al, 2020), which in addition to slope and intercept returns Pearson R and regression p-value. Clustering of phenotypes based on a matrix of correlation coefficients was performed using Scipy’s “scipy.cluster.hierarchy” module: the “linkage” method was used with the “weighted” option, invoking hierarchical WPGMA clustering.

## Results

### DFI values correlated with MFI and chronological age

Data collected for our study included 587 measurements of MFI, taken across 213 mice, aged from 252 to 938 days (data supplied in Supplemental Table 2). As has been generally reported for frailty indexes, MFI correlated positively with chronological age (Fig 2A; Pearson R = 0.27; p-value = 1.7 * 10^−11^). This correlation held for both females (348 measurements from 128 animals; Pearson R = 0.37; p-value = 3.8 * 10^−13^) and males (239 measurements from 85 animals; Pearson R = 0.23; p-value = 2.7 * 10^−4^).

**Figure 2:**
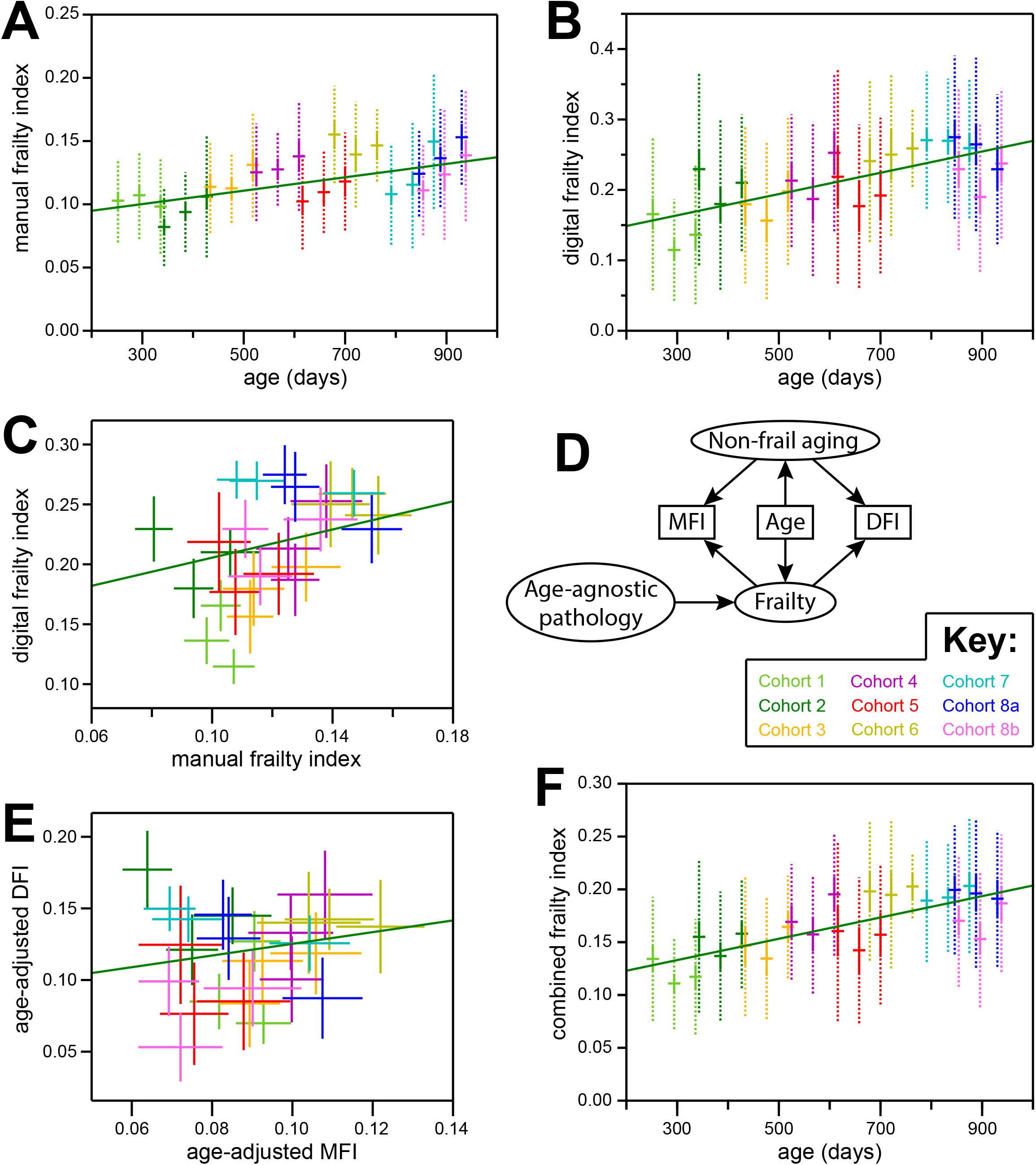
DFI positively correlated with MFI and chronological age. A) Chronological age (x-axis) versus MFI (y-axis). Mean (horizontal bars), standard errors (solid vertical bars), and standard deviations (dotted vertical bars) are shown for each of three measurement instances taken for each of eight birth cohorts (with cohort 8 split into two sub-groups: see Key). The linear regression of age versus MFI for all individual values is shown in green. B) Chronological age (x-axis) versus DFI (y-axis). Plotted as in (A). C) MFI (x-axis) versus DFI (y-axis). For each of three MFI/DFI measurements taken for each cohort, a cross is drawn whose center is the mean value on each axis for one cohort/measurement, and whose bars indicate the standard errors of those means along each axis. Colored according to the Key. The linear regression of MFI versus DFI for all individual values is shown in green. D) A path diagram depicting a model for consideration of the possible relationships between MFI, DFI, and chronological age. See Results and Discussion. E) Age-normalized MFI (x-axis) versus age-normalized DFI (y-axis), plotted as in (D). F) Chronological age (x-axis) versus combined frailty index (CFI: the average of MFI and DFI; y-axis). Plotted as in (A).

Our study also included 576 instances of measured DFI (data supplied in Supplemental Table 3). Similar to MFI, DFI correlated positively with chronological age (Fig 2B; Pearson R = 0.29; p-value = 8.0 * 10^−13^). Again, this relationship held for both females (341 measurements from 136 animals; Pearson R = 0.30; p-value = 1.4 * 10^−8^) and males (235 measurements from 88 animals; Pearson R = 0.26; p-value = 4.6 * 10^−5^).

Our study included 567 instances for which a DFI measurement was taken within seven days of an MFI measurement of the same mouse (Supplemental Table 6). DFI values were positively correlated with MFI values (Fig. 2C; Pearson R = 0.22; p-value = 1.8 * 10^−7^). This correlation held for both females (337 MFI/DFI pairs from 126 animals; Pearson R = 0.29; p-value = 8.2 * 10^−8^) and males (230 MFI/DFI pairs from 84 animals; Pearson R = 0.19; p-value = 3.1 * 10^−3^).

The correlation of both MFI and DFI with chronological age (Fig. 2C) implied that the correlation between MFI and DFI could have been mediated by chronological age rather than the intended mediator, physical frailty (Fig. 2D). To evaluate that possibility, the residuals of both MFI and DFI from the regression of each with chronological age were compared. Though weakened versus the raw values, the correlation between these residuals remained positive and statistically significant (Fig. 2E; Pearson R = 0.15; p-value = 2.7 * 10^−4^).

The significance of correlation between age-adjusted MFI and age-adjusted DFI provided reassurance that these two metrics did indeed reflect overlapping aspects of physiological decline. Furthermore, the low values of correlation coefficients amongst these two metrics and chronological age were expected given the stochastic nature of the underlying phenotypes. Nonetheless, each metric was likely to have captured aspects of age-related physiological decline missed by the other. To assess the collective potential for improvement of both metrics, we averaged their outputs into a combined frailty index and evaluated it versus chronological age. The extent of correlation was greater (Pearson R = 0.34) and more significant (p-value = 3.5 * 10^−17^) than for either individual score (Fig. 2F), suggesting opportunities for further improvement in this field.

### Individual DFI components provided variable utility in the test cohort

Our implementation of DFI included eight measurements, reflecting four arenas of function: physical fitness capability, environmental engagement, circadian rhythm, and body condition (two measurements per arena; see Methods). In all cases, methods were developed and parameterized using separately-collected training data, and the relevances to chronological age and MFI in our large test cohort varied considerably (Fig. 3A). Generally, the gait/wheel and circadian components of the DFI were correlated with both MFI and chronological age, though to varying degrees. Meanwhile, components designed to detect frailty based on nesting material, coat quality, and body weight change shared little correlation with those benchmarks.

**Figure 3:**
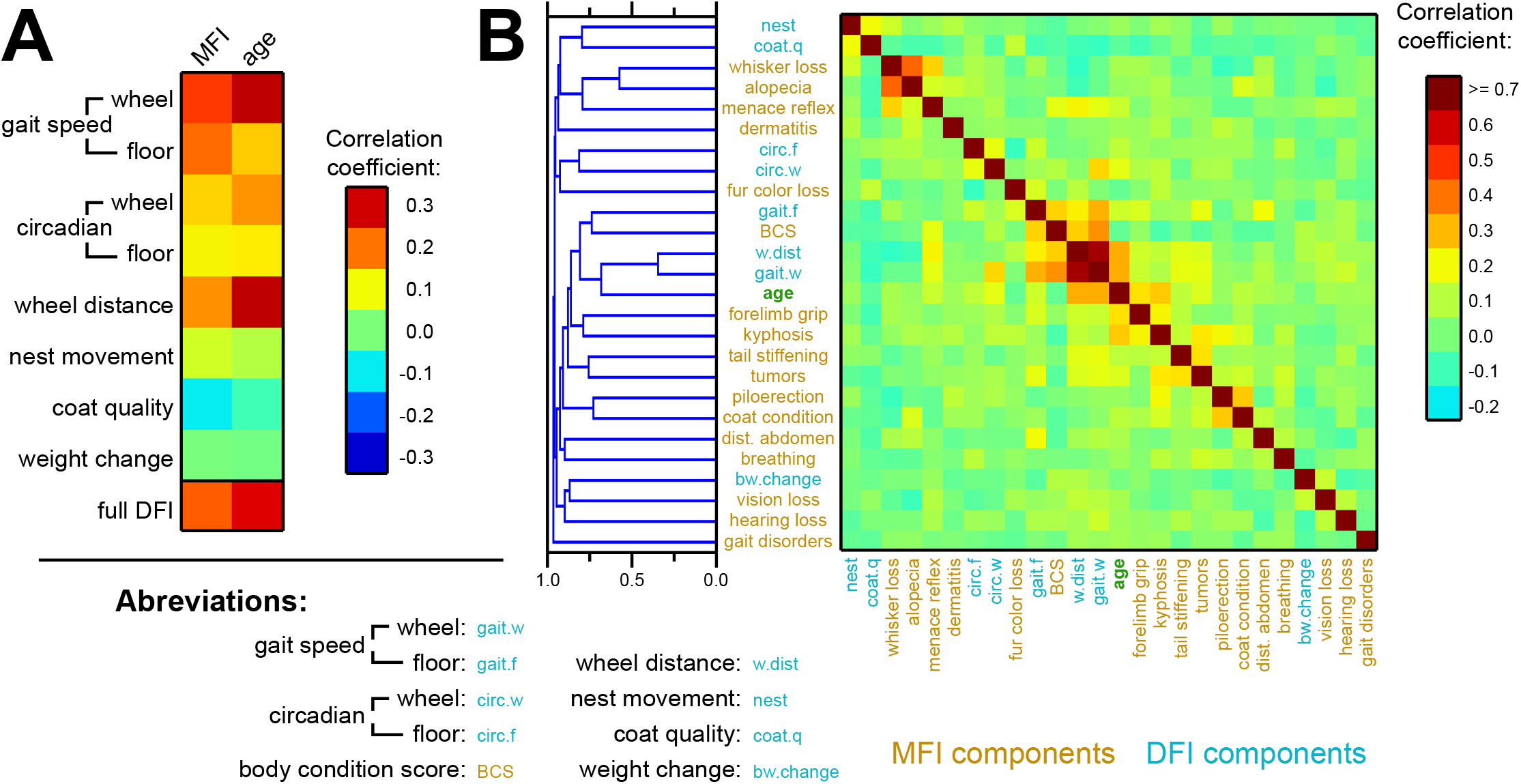
Most DFI components positively correlated with MFI and chronological age. A) A heat map indicating the correlation coefficients between individual components of the DFI (and, at bottom, the full DFI) versus MFI and chronological age. B) Hierarchical clustering (left) of individual MFI and DFI components, and chronological age, based on the correlation coefficients depicted in the heatmap (right). MFI components are labeled in orange; DFI components are labeled in blue; chronological age is labeled in green. Only MFI components observed with non-zero values >5 times across the study are included.

The clustering of DFI and MFI components based on the all-by-all correlation matrix revealed little structure (Fig. 3B). Few instances of substantial correlation between frailty traits were observed. Versus either the higher-order parameters (MFI, chronological age) or the component parameters of MFI with seeming relevance, multiple explanations may account for missing correlation aside from true irrelevance of the traits. Among them: poor model performance at measuring phenotypes; inappropriate parameterization of frailty thresholds; or sparsity of instances of certain types of frailty in a cohort of mostly healthy mice. Below, we discuss the derivation and performance of each DFI parameter.

### Walking and running statistics robustly captured age-related decline

Voluntary wheel-running is an activity known to be influenced by the physical health of mice (Greenwood & Fleshner, 2019). Our capture of elapsed time for each spin of the wheel (see Methods) permitted multiple facets of health and behavior to be probed using those data. First, the total amount of wheel running could be determined, measured as total number of spins per day. This parameter captured an aspect of environment engagement (quantity of interaction with an enrichment item) while also presumably reflecting physical endurance and athletic performance. In both our C57BL/6J and J:DO training sets, this feature declined sharply with chronological age, appearing to approach zero asymptotically as mice became old (Fig. 4A). When parameterized to frailty, strong correlation with chronological age was maintained (Fig. 4B; R = 0.32, p-value = 1.4 * 10^−14^), and correlation with MFI was achieved (R = 0.18, p = 1.8 * 10^−5^).

**Figure 4:**
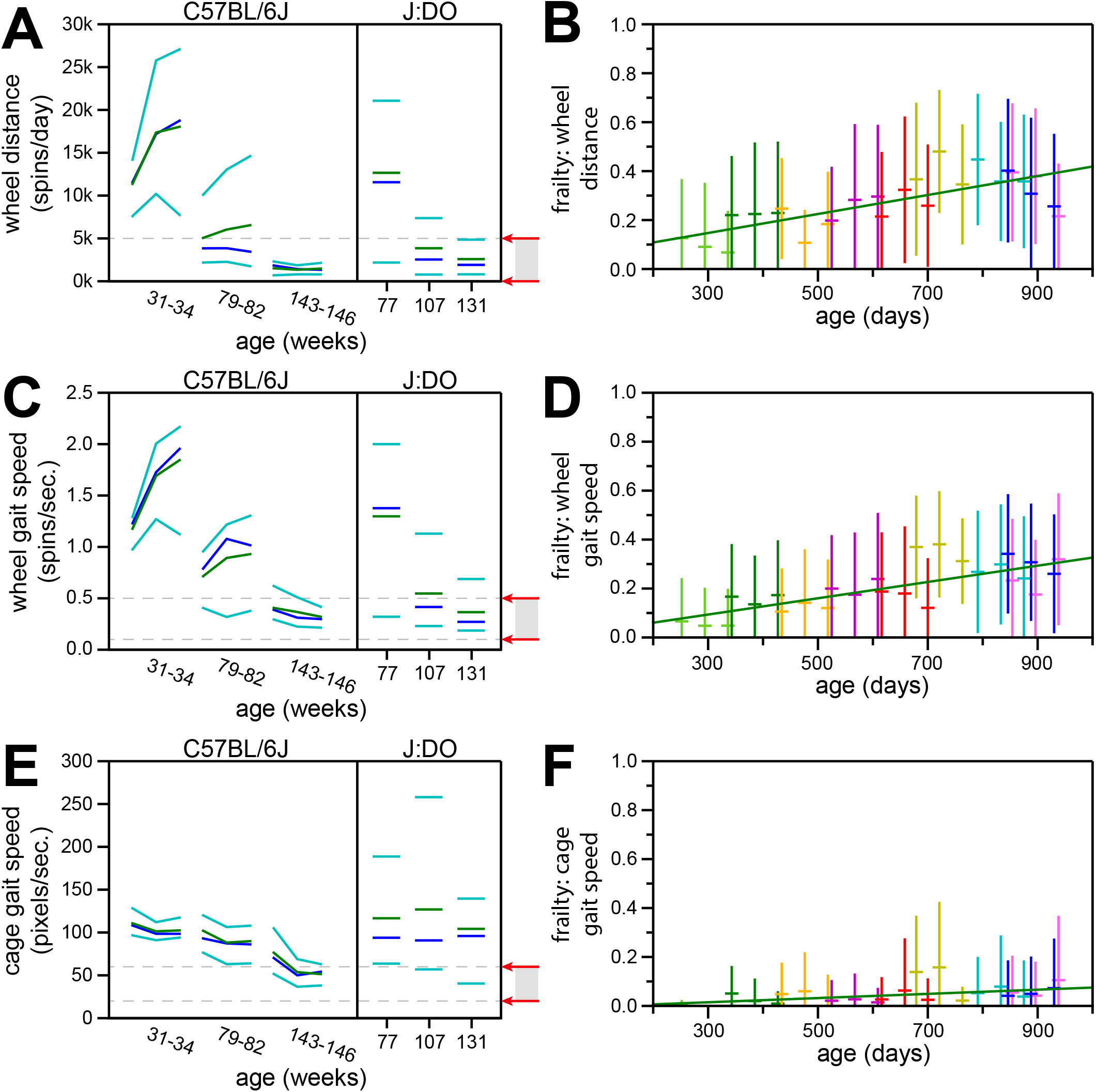
DFI components based on ambulatory activity captured age-related decline. A) Training set distributions and frailty parameterization for total wheel distance. Left: statistics for three consecutive, week-long measurements each, for three cohorts of differently-aged C57BL/6J mice. Right: similar statistics for single-week measurements from differently-aged J:DO mice. For each cohort, green bars indicate the mean value; dark blue bars indicate the median value; and light blue bars indicate the 10th and 90th percentiles. Far right: red arrows indicate the threshold values for a frailty score of zero (upper arrow) or one (lower value). Frailty values in between the thresholds (gray rectangle) were given linearly intermediate frailty values (see Methods). B) Chronological age (x-axis) versus wheel distance frailty scores (y-axis) for the main study population. Mean (horizontal bars) and standard deviations (vertical bars) are shown for each of three measurements taken for each of eight birth cohorts (with cohort 8 split into two sub-groups: see Key from Fig. 2). The linear regression of age versus wheel distance frailty score, for all individual values, is shown in green. C) Training set distributions and frailty parameterization for wheel gait speed. Plotted as in (A). D) Chronological age (x-axis) versus wheel gait speed frailty scores (y-axis) for the main study population. Plotted as in (B). E) Training set distributions and frailty parameterization for cage-floor gait speed. Plotted as in (A). F) Chronological age (x-axis) versus cage-floor gait speed frailty scores (y-axis) for the main study population. Plotted as in (B).

Gait speed on the wheel, interpreted in terms of athletic performance, was also derived from the wheel rotation frequencies (see Methods). Like total wheel-running, this parameter declined sharply with chronological age in both C57BL/6J and J:DO training-set mice (Fig. 4C). This relationship was maintained when parameterized to frailty (Fig. 4D; R = 0.32; p-value = 8.9 * 10^−15^), and correlation was achieved with MFI (R = 0.24; p-value = 3.3 * 10^−9^). Gait speed on the wheel has a lower-intensity counterpart in ambulation around the floor of the cage. The context is different (slower ambulation in the context of everyday activity, versus purposeful exercise-based enrichment), but the relationships with chronological age and MFI were similar. A strong decline with age among both C57BL/6J and J:DO training-set mice (Fig. 4E) was maintained when parameterized for frailty (Fig. 4F; R = 0.14; p-value = 8.6 * 10^−4^) and was also correlated with MFI (R = 0.21; p-value = 6.1 * 10^−7^).

### Circadian regulation of movement decreased with age

The multi-day, continuous-monitoring nature of our video data for DFI allowed us to uniquely incorporate circadian rhythm into our assessment of frailty. We implemented a measure of circadian behavior that contrasted the amount of observed movement between alternating twelve-hour blocks. This intentionally coincided with the alternating 12-hour light/dark periods experienced by the mice (see Methods), but we did not require strict adherence to the light cue. Rather, our implementation measured the overall periodicity of behavior. There are other methods for evaluating circadian rhythms (Shimomura, 2001), but this one captures a known pathology of aging: the maintenance of activity during what should be a rest period (Musiek & Holtzman, 2016).

Circadian regulation of ambulatory activity was observed to lessen with age among both C57BL/6J and J:DO training-set mice, both on the wheel (Fig. 5A) and on the floor of the cage (Fig. 5B). When parameterized for frailty, the decline of circadian activity with age was maintained, both for wheel-running activity (Fig. 5C; R = 0.18; p-value = 2.1 * 10^−5^) and for movement on the floor of the cage (Fig. 5D; R = 0.11; p = 6.9 *10^−3^). Both metrics also correlated with overall MFI, with correlation coefficients of 0.13 (p-value = 2.1 * 10^−3^) and 0.11 (p-value = 0.011) for wheel-based and cage floor-based activity, respectively.

**Figure 5:**
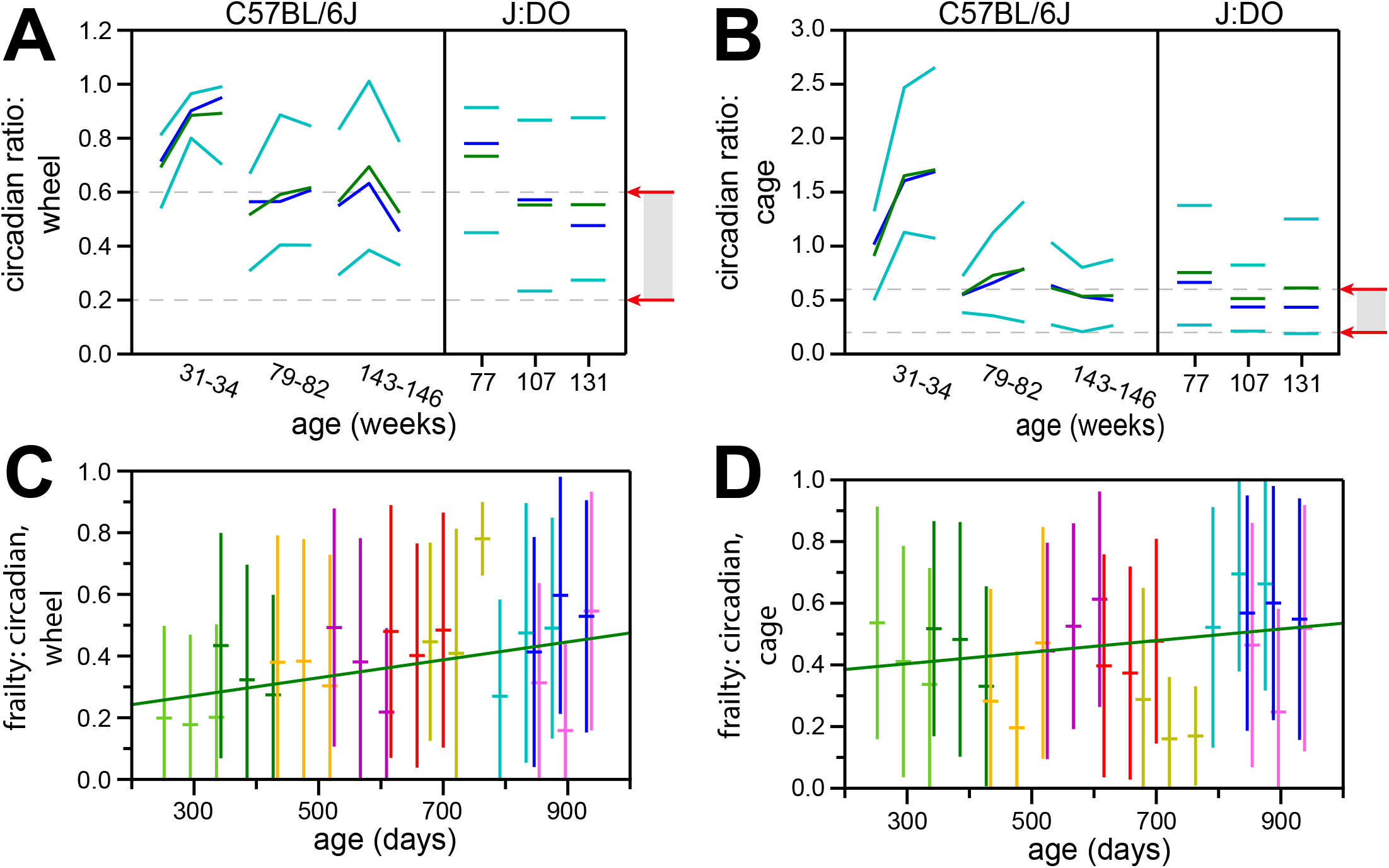
DFI components based on circadian activity captured age-related decline. A) Training set distributions and frailty parameterization for the circadian ratio of wheel-running activity (see Methods). Plotted as described for Fig. 4A. B) Training set distributions and frailty parameterization for the circadian ratio of movement on the floor of the cage (see Methods). Plotted as described for Figure 4A. C) Chronological age (x-axis) versus circadian wheel-running frailty scores (y-axis) for the main study population. Plotted as in Fig. 4B. D) Chronological age (x-axis) versus frailty scores for circadian movement on the floor of the cage (y-axis) for the main study population. Plotted as in Fig. 4B.

### Frailty measurements of coat quality, body-weight change, and nest movement showed no statistical change with age

Three components of our DFI implementation seemed justifiable to include but nonetheless performed poorly against the chronological age and/or MFI values of our test set. The first of these sought to quantify the movement of nest material. It is known that mice in pain or distress will expend less effort constructing nests (Gaskill et al, 2013), a phenomenon that is most effectively observed when mice are first provided with nesting material (Giménez-Llort & Torres-Lista, 2021). Our DFI analysis was not timed to the specific time at which the mouse was introduced to a new cage with new nest material and was therefore limited to the continued maintenance and modification of existing nests. Frailty was therefore parameterized to only capture instances of severely depleted engagement with the nest and was therefore only sparsely observed (Fig. 6A). Though nest-moving frailty trended positively with both chronological age (R = 0.05) and MFI (R = 0.07), in neither case was the association statistically significant. This parameter had no obviously analogous component of MFI. The most highly correlated MFI component was “whisker loss” (R = 0.14; p-value = 9.3 * 10^−4^).

**Figure 6:**
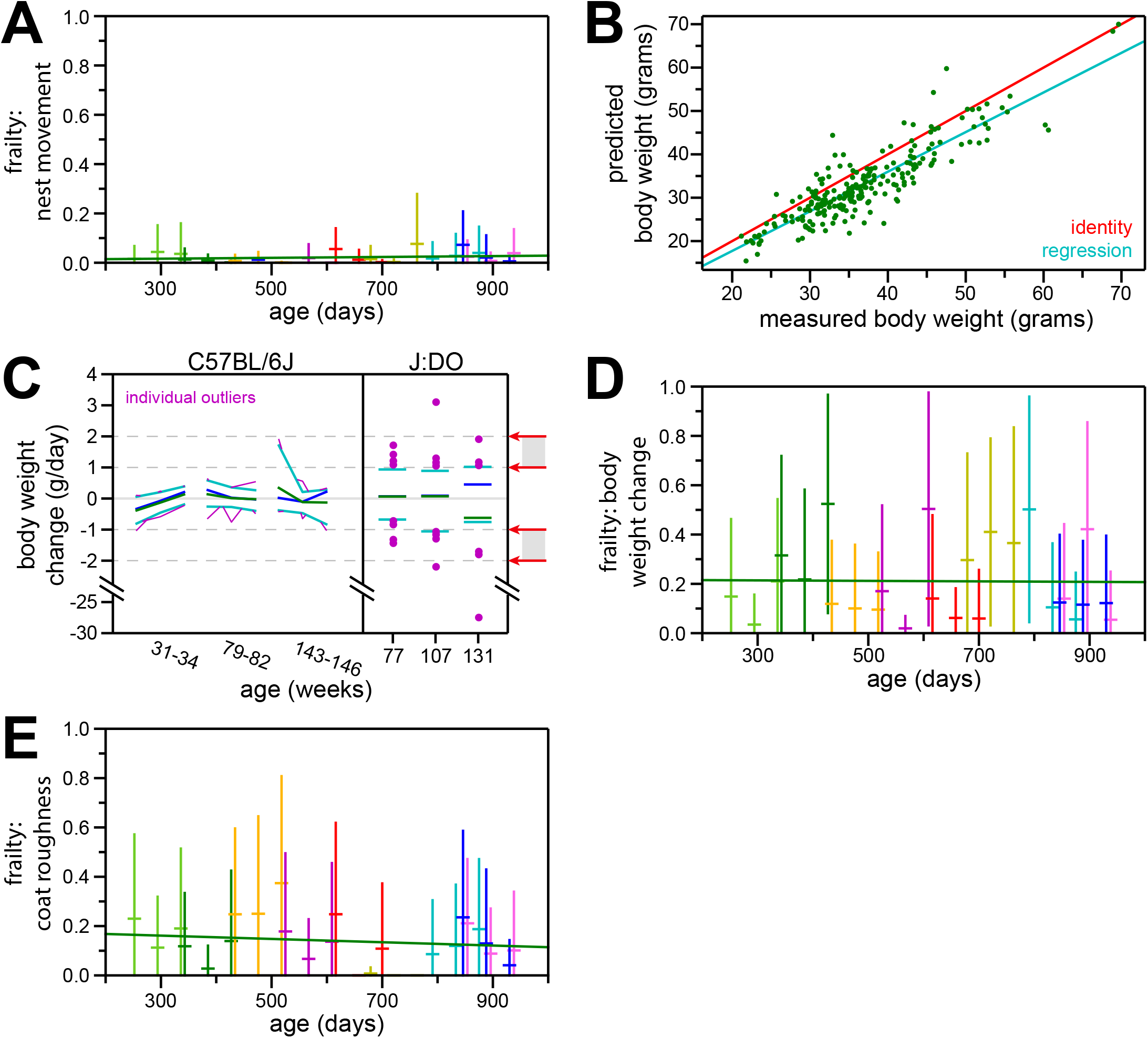
DFI components based on nest movement, rapid body weight change, and coat roughness failed to capture age-related decline. A) Chronological age (x-axis) versus nest movement frailty scores (y-axis) for the main study population. Plotted as in Fig. 4B. B) Performance of the body weight prediction model using 24-hour averages. Measured body weights (x-axis) versus predicted body weights based on statistical averaging of model outputs from across the same 24-hour period (y-axis) are plotted in green: each point is a measurement/prediction instance. Shown along with the calculated regression line (blue) and identity line (red). C) Training set distributions and frailty parameterization for changes to body weight. Plotted as described for Fig. 4A, but with individual values outside of the 10th and 90th percentiles additionally plotted (purple), and with both positive and negative thresholds for the absolute-value-based frailty score depicted (inner thresholds correspond to zero; outer thresholds correspond to one; see Methods). D) Chronological age (x-axis) versus frailty scores based on changes in body weight (y-axis) for the main study population. Plotted as in Fig. 4B. E) Chronological age (x-axis) versus frailty scores based on coat roughness (y-axis) for the main study population. Plotted as in Fig. 4B.

The second poorly-performing DFI component was “body weight delta”. The foundation of this parameter was a body-weight prediction tool. First developed and tested on an independent data set to predict body weight for a mouse by averaging from estimates taken across a day’s worth of video (sampled every ten minutes; results averaged across 144 estimates/day), this model produced daily estimates within ∼4 grams of the laboratory measured body weights (R = 0.88 across a test-set of 250 measurements with an SD = 33.9 grams; p-value = 7.2 * 10^−77^; Fig. 6B). For frailty, this prediction tool was used to identify rapid gain or loss of weight by regressing across estimates taken once-per-minute across each designated DFI measurement period

Rapid weight gain or loss is a known harbinger of mortality in mice that is often considered as a humane endpoint (Toth, 1997). It is sometimes included in traditional frailty assessments, but doing so requires the manual collection of weights at disparate time points, therefore modifying the nature of the frailty index into a longitudinal assessment (e.g. Mach et al, 2022). For DFI, the trend in body weight across the ∼one-week interval of observation time was used to calculate body weight dynamics as a single-observation measurement. Among both C57BL/6J and J:DO training-set mice, this method identified body weights as being highly stable, with a limited number of outlier exceptions (Fig. 6C). For frailty, this metric was parameterized to capture those outliers. The resulting frailty scores were significantly correlated with neither chronological age (Fig. 6D; R = -0.01; p-value = 0.89) nor MFI (R = -0.00; p-value = 1.00). The two most highly correlated individual MFI components were “body condition score” (R = 0.09; p-value = 0.025) and “vision loss” (R = 0.13; p-value = 2.2 * 10^−3^). The correlation with “body condition score” suggested that some appropriate signal was captured by this metric, but that signal was overwhelmed by the noise of the measurement. Reassuringly, MFI “body condition score” had a negative, non-significant correlation with chronological age in our test cohort (R = -0.04; p-value = 0.30), suggesting that the inability of this parameter to correlate with chronological age was not entirely due to inaccuracy of the measurement.

The third poorly-performing DFI component was “coat quality”. For this metric, the Sobel gradient estimator was used to quantify the roughness of the portion of images containing the mouse, as determined by a segmentation model (see Methods). Parameterized for frailty, no correlation was found with chronological age (Fig. XE; R = -0.06; p-value = 0.19), and a negative correlation was found with MFI (R = -0.10; p-value = 0.018). Conceptually, “coat quality” could have encompassed multiple components of the MFI: “alopecia”, “loss of fur color”, “dermatitis”, “coat condition”, and/or “piloerection”. All of these were expected to increase the visual unevenness of the mouse’s coat. However, only one of these five MFI components positively correlated with DFI “coat quality”, and it was the only one of the five with a statistically significant correlation: “loss of fur color” (R = 0.13; p-value = 2.7 * 10^−3^). That MFI value also had a negative correlation with chronological age in our frailty study, though that correlation was not statistically significant (R = -0.04; p-value = 0.32). Taken together, these results suggested that the Sobel estimate accurately captured the color-loss aspect of coat condition, and that supplementation with additional measurements to better capture other aspects of coat condition would be preferential to abandonment of this parameter.

## Discussion

Here, we implemented a digital frailty index (DFI) for mice based entirely on the computational analysis of continuously-collected home-cage video footage. We evaluated performance of our DFI against both chronological age and manual frailty index - MFI - and found it to correlate with both metrics in a cross-sectional study using J:DO mice. Importantly, the latter experimental design condition (the use of genetically diverse mice, with varying builds and coat colors) demonstrated our method to not be dependent for its effectiveness or relevance to a particular inbred strain and therefore of potential use across a wide range of in vivo models.

Frailty indexes are so named because they are meant to reflect a lack of physical well-being. They are not explicitly designed to be predictors of chronological age, but the natural physiological decline that accompanies age results in a natural and expected age-dependence. Traditional frailty indexes are not strategically designed to capture chronological age aside from its effect on disability. However, it has been shown that even the components of a traditional mouse frailty index can be weighted to optimize the accurate prediction of age (Schultz et al, 2020). The correlation between DFI and MFI across an age-diverse cohort therefore did not guarantee that DFI was actually performing as an index of frailty, but rather could have been performing as a chronological age predictor (FIG. 2D). To allay that possibility, we evaluated MFI versus DFI scores after regressing the age component out of each, observing the correlation between the remaining age-independent frailty residuals to retain their correlation (Fig. 2E).

Home-cage DFI represents a unique practical step forward for longitudinal health assessments in terms of vivarium operational efficiency and scalability. Since observations are taken in a home-cage environment, highly-trained personnel are not required to take measurements, just the vivarium husbandry staff required for animal feeding and care. Furthermore, in order to be meaningful, traditional frailty assessment requires consistent scoring across large and longitudinal. In practice, substantial interobserver variability drives variance up as studies are scaled up and additional researchers are required to collect data, negating much of the statistical power that would otherwise accompany a larger data set. On the other hand, DFI can be applied in a uniform manner regardless of the experimental scale. This advantage also applies to the length of studies: for long-term aging studies, it can be difficult to guarantee the consistent staffing required for meaningful longitudinal frailty data.

Home-cage DFI also represents a unique step forward for longitudinal health assessments in terms of improving animal welfare. The home-cage context of the measurement eliminates a substantial but otherwise almost unavoidable source of stress for the animals: handling. Frailty assessments are generally non-invasive and therefore already considerate of animal welfare, but animal handling remains and is difficult to avoid. The technical success of video-based DFI, itself a multi-faceted measurement, demonstrates the utility of video as a physiological measurement tool that can abrogate the need for animal handling.

While the DFI implementation presented here substantially advances the operational and animal welfare aspects of longitudinal health studies and meets the correlative requirements for general use, it certainly also leaves room for improvement. It will likely be refined and augmented in the future, to provide greater sensitivity and nuance to the detection of animal frailty. Nonetheless, even in its current state, it serves as a proof of principle for the aspirational concept of an operationally efficient and humane digital vivarium (Baran et al, 2022).

## Data & code availability

Code for running DFI analysis on a set of videos is available at https://github.com/graham-calico/DFI_v1.

Supplemental Table 1: Census data on all mice obtained for this study.

Supplemental Table 2: MFI score data.

Supplemental Table 3: DFI score data: overall score and parameterized components.

Supplemental Table 4: pre-parameterized values for DFI components.

Supplemental Table 5: time intervals of DFI video collection, compliant with DFI software.

Supplemental Table 6: qualifying MFI/DFI pairs.

## Acknowledgements

This study was funded by Calico Life Sciences LLC.

J.G.R. and E.M.K. conceptualized the project. J.G.R., E.M.K. and A.D.F. contributed to study design. V.K., P.C., and E.M.K. managed project organization and activities. R.K., O.W., S.S., W.L., N.V., S.C., C.A.S., and C.L. performed digital data acquisition and organization. P.Y., A.D.F., J.Z-S., J.K., M.C., and P.C. contributed to the acquisition of animal data and observations. J.G.R. performed data analysis and wrote the manuscript. All authors reviewed the manuscript and approved the final version of the manuscript for submission.

